# Enhancing associative learning in rats with a computationally designed training protocol

**DOI:** 10.1101/2022.06.08.495364

**Authors:** Xu O. Zhang, Yili Zhang, Claire E. Cho, Douglas S. Engelke, Paul Smolen, John H Byrne, Fabricio H. Do-Monte

## Abstract

**Background:** Learning requires the activation of protein kinases with distinct temporal dynamics. In *Aplysia,* nonassociative learning can be enhanced by a computationally designed learning protocol with intertrial intervals (ITIs) that maximize the interaction between fast-activated protein kinase A (PKA) and slow-activated extracellular signal-regulated kinase (ERK). Whether a similar strategy can enhance associative learning in mammals is unknown.

**Methods:** We simulated 1,000 training protocols with varying ITIs to predict an optimal protocol based on empirical data for PKA and ERK dynamics in rat hippocampus. Adult male rats received the optimal protocol or control protocols in auditory fear conditioning and fear extinction experiments. Immunohistochemistry was performed to evaluate phosphorylated cAMP responsive element binding (pCREB) protein levels in brain regions implicated in fear acquisition.

**Results:** Rats exposed to the optimal conditioning protocol with irregular ITIs exhibited impaired extinction memory acquisition within the session using a standard footshock intensity, and stronger fear memory retrieval and spontaneous recovery with a weaker footshock intensity, compared to rats that received massed or spaced conditioning protocols with fixed ITIs. Rats exposed to the optimal extinction protocol displayed improved extinction of contextual fear memory and reduced spontaneous recovery compared to rats that received standard extinction protocols. Moreover, the optimal conditioning protocol increased pCREB levels in the dentate gyrus (DG) of the dorsal hippocampus, suggesting enhanced induction of long-term potentiation.

**Conclusion:** These findings demonstrate that a computational model-driven behavioral intervention can enhance associative learning in mammals, and may provide insight into strategies to improve cognition in humans.

## INTRODUCTION

Long-term memory (LTM) formation is believed to be mediated by synaptic plasticity including long-term potentiation (LTP) or its invertebrate analogue long-term facilitation (LTF), which require gene expression and protein synthesis (1–5). Studies over decades have investigated LTP/LTF and their underlying molecular processes as potential targets to enhance learning or restore memory deficits in laboratory animals. However, interventions using systemic cognitive enhancers or intracerebral pharmacological manipulations (6–10) are either based on trial-and-error approaches in specific model systems or are highly invasive, preventing or hindering translation to humans.

An alternative approach to enhance learning and memory is to develop computational models to predict optimal learning protocols based on dynamics of intracellular molecular cascades that underlie LTM formation and LTP/LTF induction (11–13). Studies have identified activation of PKA (14–16) and of the mitogen-activated protein kinase (MAPK) isoform ERK (17–19) as essential cascades for LTF. These two pathways converge to phosphorylate transcriptional factors such as CREB, which subsequently induce plasticity-related genes during LTF induction (20–24). These two pathways exhibit distinct kinetics of activation (13, 16, 25), suggesting that the temporal activity patterns, and activation overlap, of these pathways may constitute an important target to enhance associative learning. We previously demonstrated that a computationally designed protocol with irregular ITIs, predicted to maximize overlap of PKA and ERK activities, enhances LTF and nonassociative learning, specifically long-term sensitization of the tail elicited siphon-withdrawal reflex, in *Aplysia* (26).

Substantial similarities between molecular processes of LTF in invertebrates and LTP in mammals make it plausible that strategies used to enhance LTF and nonassociative learning in invertebrates could enhance LTP and associative learning in mammals. PKA activation in rodent hippocampus and amygdala is required for LTP and LTM (27, 28). Activation of ERK/MAPK cascades and their cross-talk with PKA are required for CREB phosphorylation, induction of plasticity-related genes, and LTP induction (29–32). Although the dynamics of PKA and ERK activation differ between invertebrates and mammals (33–35), and may vary across different brain subregions and behavioral protocols in rodents, the chronological order of activation for these two intracellular molecules is evolutionarily-conserved across species. In fact, a PKA peak precedes the nuclear translocation of ERK and the subsequent activation of CREB in Aplysia (16, 25), mice (29), and rats (36–38) which provides strong support for our model.

Therefore, we tested if our invertebrate LTF model (26) can be adapted to computationally design an optimal associative learning protocol in mammals. Because the majority of the above literature on LTP induction has focused on fear conditioning experiments, we employed this well-established paradigm to investigate associative learning (39) and used extant empirical data to model PKA and ERK dynamics in rat hippocampus (33–35), a critical brain region implicated in the formation of associative memories (40–42). We simulated 1,000 different training protocols with varying ITIs to identify an optimal protocol with irregular ITIs that maximizes the overlap of PKA and ERK activities, thereby predicting the formation of stronger memory than training protocols with the same number of stimuli and fixed ITIs. We then tested this optimal protocol in auditory fear conditioning and fear extinction paradigms in rats.

## RESULTS

### A computational model based on PKA and ERK dynamics identified an optimal fear conditioning protocol

The model in (26) was adapted to simulate PKA and ERK dynamics in rat hippocampus during LTP induction (**Fig. 1A**). To the best of our knowledge, hippocampus is the only rat brain region in which ERK dynamics has been measured with a temporal resolution of minutes following a stimulus. We simulated fear conditioning, in which the pairing of a conditioned stimulus (CS) with an unconditioned stimulus (US) is represented by *Stim*. The conditioned responses are assumed to be proportional to the peak value of *inducer*. A single CS-US pairing produced little overlap between PKA and ERK pathways (**Fig. 1B 1**). We then simulated 1,000 fear conditioning protocols, with 4 trials and both fixed and irregular ITIs. These simulations included a Short Conditioning (SC) protocol with 4 trials and a fixed ITI of 270 s. As a reference control, we also simulated a Regular Conditioning (RC) protocol with 8 trials and the same ITI of 270 s. These protocols respectively resemble previous protocols used for short and regular fear conditioning in rats (43, 44). Simulations identified an optimal conditioning protocol with 4 trials and irregular ITIs of 8, 8, and 16 min (**Fig. 1B 2-1B4**), termed Optimal Short Conditioning (OSC). PKA activity induced by the last trial of OSC had a much larger overlap with the pERK curve than did PKA induced by SC. Therefore, OSC induced a much higher peak level of *inducer* (red). We therefore predicted OSC would produce stronger long-term memory in rats than standard SC.

**Figure 1.**
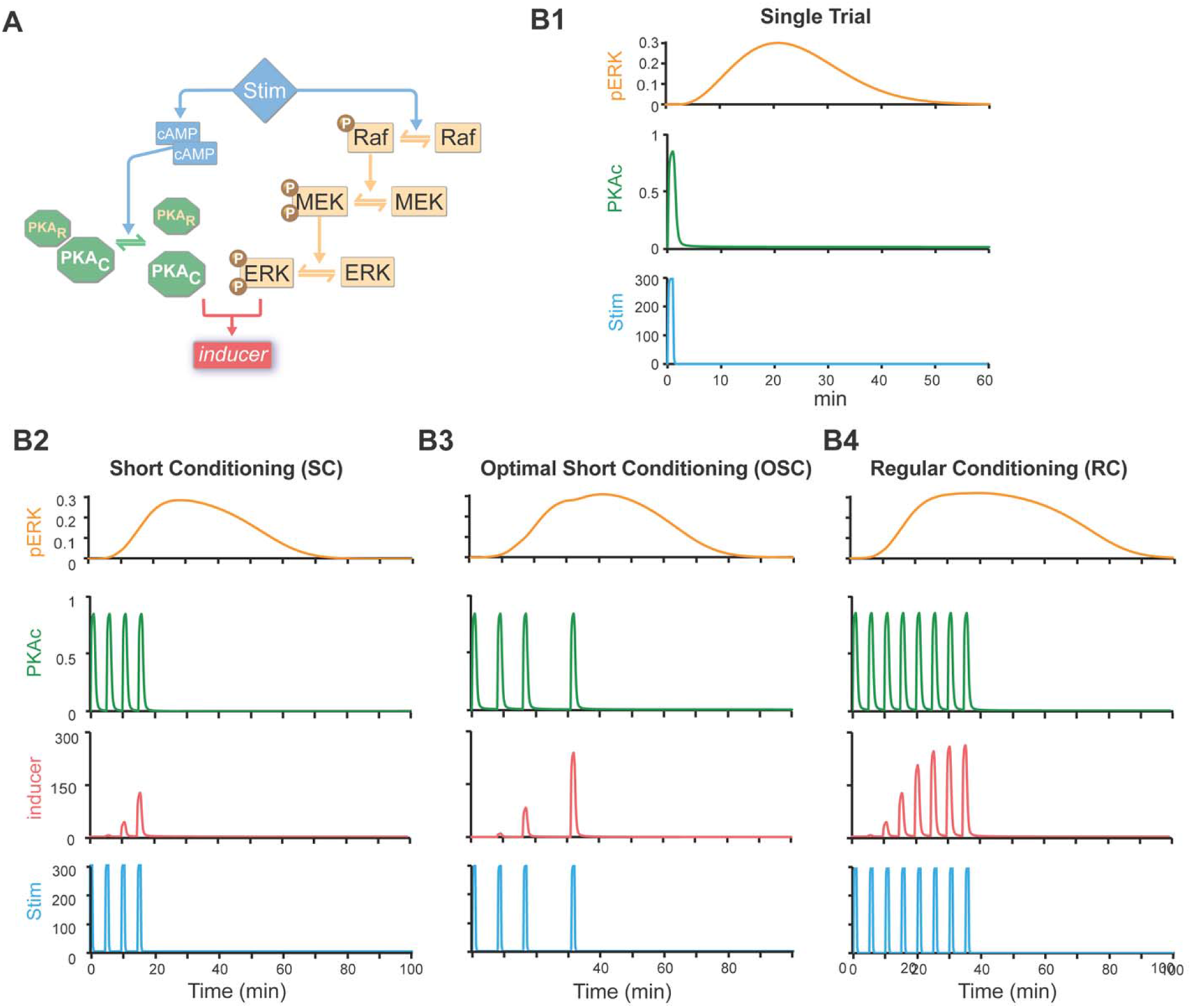
Computational simulations of PKA and ERK pathways predict an optimal protocol for fear conditioning. **A**, Schematic of the model. Stimulus (*Stim*, μM) rapidly activates PKA via cAMP pathway, and activates ERK more slowly via Raf-MEK pthway. The synergistic interaction between PKA and ERK pathways is quantified by a variable *inducer*, which corresponds to the efficacy of the stimulus in inducing LTP. ERK kinetics were described by differential equations (see Supplementary Methods) with parameter values reproducing empirical findings that ERK activity reaches peak levels 15∼20 min after BDNF treatment or tetanic stimuli in rat hippocampus acute slices (35, 78). Equations describing PKA kinetics simulated data showing that PKA is transiently activated within 2 min after LTP induction in slices from rat hippocampus or within 5 min *in vivo* after spatial discrimination task training (33, 34). **B1**, Simulated time courses of activated ERK (pERK, μM) and activated PKA (PKAc, μM) in response to one trial of *Stim*. **B2**, Simulated time courses of pERK, PKAc, and *inducer* in response to a 4-trial protocol with regular ITIs of 4.5 min (short conditioning, SC). **B3**, Simulated time courses of pERK, PKAc, and *inducer* (nM) in response to a 4-trial protocol with computationally designed intervals (optimal short conditioning, OSC). **B4**, Simulated time courses of pERK, PKAc, and *inducer* (nM) in response to an 8-trial protocol with regular ITIs of 4.5 min (regular conditioning, RC).

### The optimal short conditioning protocol increases fear memory compared to the standard short conditioning protocol

Because the hippocampus is required for context encoding during auditory fear conditioning (45, 46), we used two different chambers (context A and context B) to assess the contribution of the context to the auditory fear memory(**Fig. 2A**). Our pilot experiments and a series of recent fear conditioning studies have shown that female rats exhibit higher active defensive responses (i.e., darting) and lower passive defensive responses (i.e., freezing) than male rats (47–50). We therefore performed our experiments in males which exhibit enhanced freezing behavior, the main index of fear memory adopted in our study. On Day 0, rats were familiarized with context A for 20 min. They were pre-assigned to Short Conditioning (SC), Optimal Short Conditioning (OSC), or Regular Conditioning (RC) groups so that baseline freezing and locomotion were similar among groups (**Table 1**). On Day 1, rats were placed into context A and exposed to one nonreinforced habituation tone followed by the SC, OSC, or RC protocols (**Fig. 2B**). All groups reached high levels of freezing (**Fig. 2C**) and reduced locomotion at the end of the fear acquisition session.

**Figure 2.**
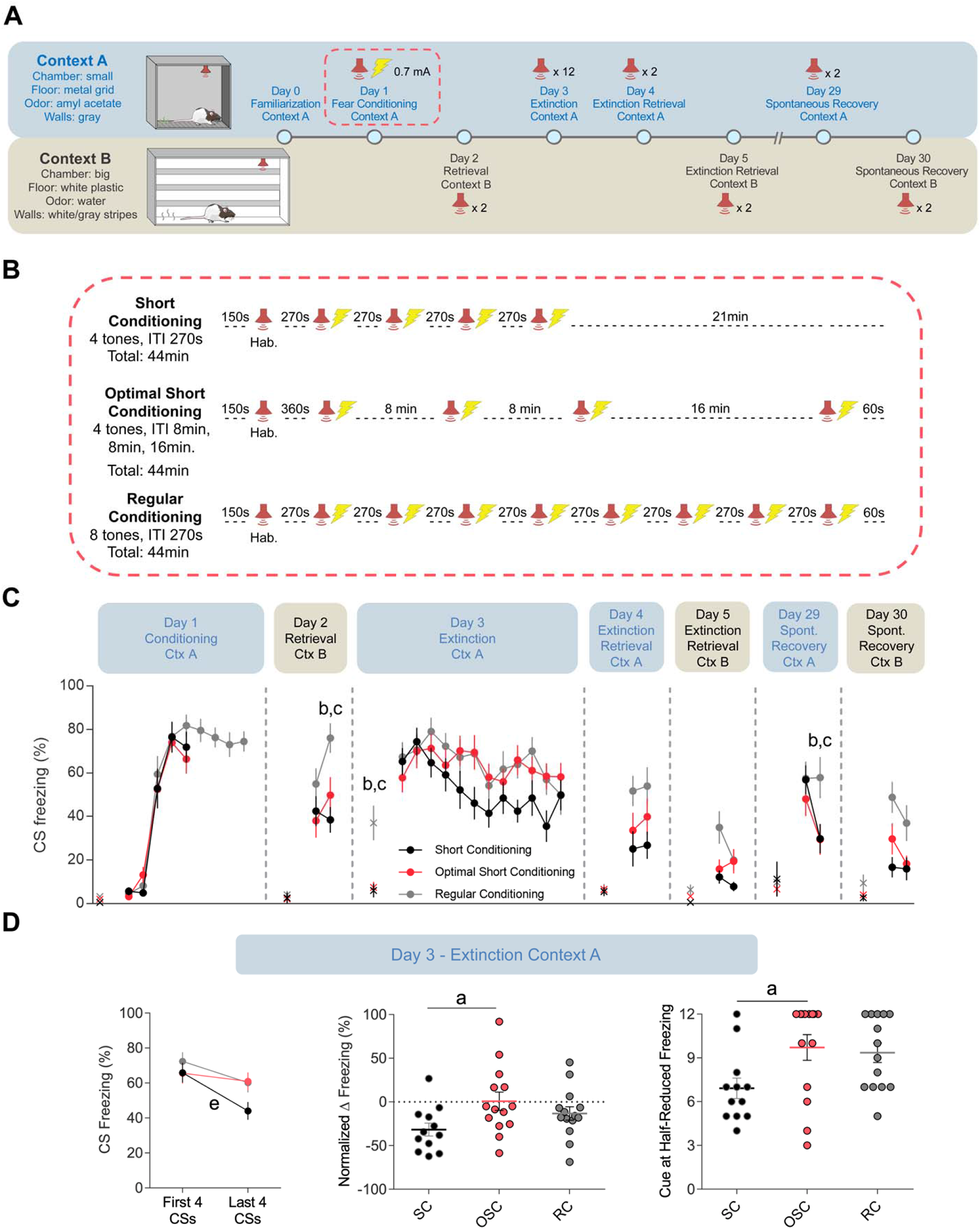
Computationally designed protocol partially enhanced fear conditioning in rats. **A,** Schematics of the enhanced fear conditioning experimental procedures. Top: tests that were conducted in Context A. Bottom: tests that were conducted in Context B. **B,** Schematics of the fear conditioning protocols for SC (n = 12), OSC (n = 14), and RC (n = 14) groups. Following a habituation (Hab.) tone, rats received multiple trials of a CS (3 kHz tone, 30 s) that co-terminated with a US (footshock, 0.7 mA, 0.5 s). **C,** Freezing levels during CS presentations of each group across the experiment. Two-way Repeated-Measure ANOVA for each day followed by Holm-Sidak’s post hoc test. Letters *a, b*, and *c* indicate pairwise post hoc tests with p < 0.05: *a*, OSC vs. SC; *b*, OSC vs. RC; *c*, SC vs. RC. ‘X’ denotes the pre-tone freezing levels. **D,** OSC group is resistant to extinction while SC group shows significantly more extinction. *Left*, freezing levels during the first 4 and last 4 CS presentations. Paired Student’s t-test. Letter *e* indicates test with p < 0.05: SC, Last 4 CS vs. First 4 CS. *Center*, normalized change of the freezing level during extinction, as indicated by the difference of the freezing levels between last 4 and first 4 CS presentations as a percentage of the freezing level of the first 4 CS presentations ((Last 4 CS – First 4 CS)/First 4 CS). One-way ANOVA followed by Tukey’s post hoc test. Letter *a* indicates pairwise test with p < 0.05: OSC vs. SC. *Right*, the number of cues required to reach to a 50% reduction of the original freezing level (average of freezing level during the first two cues). Kruskal-Wallis’s test followed by Dunn’s multiple comparisons test. Letter *a* indicates pairwise test with p < 0.05: OSC vs. SC. Data shown here and in the next illustrations as mean ± SEM.

**Table 1.** Statistical Analyses Results

| Figure | Experiment Day | Description | Statistical test | Omnibus test | Omnibus p value | Post-hoc test | Post-hoc comparisons | Post-hoc p |
| --- | --- | --- | --- | --- | --- | --- | --- | --- |
| 2 | Day 0 | Baseline freezing | One-way ANOVA | $F(2,37) = 0.018$ | 0.982 | n/a | n/a | n/a |
| | | Baseline average speed | One-way ANOVA | $F(2,37) = 0.062$ | 0.939 | n/a | n/a | n/a |
| | | Baseline maximum speed | One-way ANOVA | $F(2,37) = 0.044$ | 0.956 | n/a | n/a | n/a |
| | Day 2 | Pre-tone freezing | Kruskal-Wallis | $H(2) = 0.887$ | 0.143 | n/a | n/a | n/a |
| | | Second CS freezing | Two-way RM ANOVA | $F(2,37) = 3.751$ | 0.033 | Holm-Sidak | OSC vs. SC<br>OSC vs. RC<br>SC vs. RC | 0.282<br>0.022<br>0.002 |
| | | Average speed | One-way ANOVA | $F(2,37) = 10.87$ | < 0.001 | Tukey | OSC vs. SC<br>OSC vs. RC<br>SC vs. RC | 0.024<br>0.131<br><0.001 |
| | Day 3 | Pre-tone freezing | Kruskal-Wallis | $H(2) = 13.93$ | < 0.001 | Dunn | OSC vs. SC<br>OSC vs. RC<br>SC vs. RC | >0.999<br>0.020<br>0.001 |
| | | First 2 CS freezing | Two-way RM ANOVA | $F(22,407) = 1.169$ | 0.271 | n/a | n/a | n/a |
| | | First 4 vs. last 4 CS freezing | Paired student's t-test | OSC: $t(13) = 0.869$<br>SC: $t(11) = 4.191$<br>RC: $t(13) = 2.153$ | 0.400<br>0.001<br>0.051 | n/a | n/a | n/a |
| | | Relative change in freezing | One-way ANOVA | $F(2,37) = 3.315$ | 0.047 | Tukey | OSC vs. SC<br>OSC vs. RC<br>SC vs. RC | 0.037<br>0.487<br>0.320 |
| | | Cue at half-reduced freezing | Kruskal-Wallis | $H(2) = 7.08$ | 0.029 | Dunn | OSC vs. SC<br>OSC vs. RC<br>SC vs. RC | 0.039<br>>0.099<br>0.095 |
| | Day 4 | Extinction retrieval CTX A | Two-way RM ANOVA | $F(2,37) = 0.097$ | 0.907 | n/a | n/a | n/a |
| | | Average CS freezing | One-way ANOVA | $F(2,37) = 3.944$ | 0.028 | Tukey | OSC vs. SC<br>OSC vs. RC<br>SC vs. RC | 0.508<br>0.508<br>0.023 |
| | Day 5 | Extinction retrieval CTX B | Two-way RM ANOVA | $F(2,37) = 2.984$ | 0.063 | n/a | n/a | n/a |
| | | Average CS freezing | One-way ANOVA | $F(2,37) = 4.067$ | 0.025 | Tukey | OSC vs. SC<br>OSC vs. RC<br>SC vs. RC | 0.437<br>0.236<br>0.020 |
| | Day 29 | Spontaneous recovery | Two-way RM ANOVA | $F(2,37) = 3.259$ | 0.049 | Holm-Sidak | OSC vs. SC<br>OSC vs. RC<br>SC vs. RC | 0.976<br>0.030<br>0.030 |
| | Day 30 | Spontaneous recovery | Two-way RM ANOVA | $F(2,37) = 0.774$ | 0.469 | n/a | n/a | n/a |
| 4 | Day 0 | Baseline freezing | Unpaired student's t-test | $t(16) = 0.988$ | 0.337 | n/a | n/a | n/a |
| | | Baseline average speed | Unpaired student's t-test | $t(16) = 0.419$ | 0.680 | n/a | n/a | n/a |
| | | Baseline maximum speed | Unpaired student's t-test | $t(16) = 0.225$ | 0.825 | n/a | n/a | n/a |
| | Day 1 | Freezing 2nd CS-US pairing | Two-way RM ANOVA | $F(4,64) = 1.550$ | 0.339 | Holm-Sidak planned comparison | OSC vs. SSC | 0.040 |
| | | Freezing 3rd CS-US pairing | Two-way RM ANOVA | $F(4,64) = 1.550$ | 0.339 | Holm-Sidak planned comparison | OSC vs. SSC | 0.019 |
| | Day 3 | First 2 CS freezing | Welch's t-test | $t(12.43) = 2.227$ | 0.045 | n/a | n/a | n/a |
| | | Freezing end of session (tone 12) | Two-way RM ANOVA | $F(11,176) = 2.997$ | 0.001 | Holm-Sidak | OSC vs. SSC | 0.987 |
| | Day 4 | Extinction retrieval CTX A | Two-way RM ANOVA | $F(1,16) = 0.349$ | 0.563 | n/a | n/a | n/a |
| | Day 5 | Extinction retrieval CTX B | Two-way RM ANOVA | $F(1,16) = 1.130$ | 0.304 | n/a | n/a | n/a |
| | Day 29 | Freezing first CS presentation | Two-way RM ANOVA | $F(1,16) = 1.809$ | 0.197 | Holm-Sidak planned comparison | OSC vs. SSC | 0.037 |
| 5 | Day 0 | Baseline freezing | One-way ANOVA | $F(2,12) = 0.970$ | 0.407 | n/a | n/a | n/a |
| | | Baseline average speed | One-way ANOVA | $F(2,12) = 3.706$ | 0.056 | n/a | n/a | n/a |
| | | Baseline maximum speed | One-way ANOVA | $F(2,12) = 3.839$ | 0.051 | n/a | n/a | n/a |
|  | Day 1 | Freezing 2nd CS-US pairing | Two-way RM ANOVA | F(8,48) = 1.976 | 0.070 | Holm-Sidak planned comparison | OSC vs. SSC<br>SSC vs. NS<br>OSC vs. NS | 0.144<br>0.029<br>0.001 |
|  |  | Freezing 3rd CS-US pairing | Two-way RM ANOVA | F(8,48) = 1.976 | 0.070 | Holm-Sidak planned comparison | OSC vs. SSC<br>SSC vs. NS<br>OSC vs. NS | 0.312<br>0.002<br><0.001 |
|  |  | Freezing 4th CS-US pairing | Two-way RM ANOVA | F(8,48) = 1.976 | 0.070 | Holm-Sidak planned comparison | OSC vs. SSC<br>SSC vs. NS<br>OSC vs. NS | 0.020<br>0.037<br><0.001 |
|  |  | Lateral amygdala immunohistochemistry | Two-way RM ANOVA | F(4,24) = 2.844 | 0.046 | Holm-Sidak | OSC vs. SSC<br>SSC vs. NS<br>OSC vs. NS | 0.838<br><0.001<br><0.001 |
|  |  | Basal amygdala immunohistochemistry | Two-way RM ANOVA | F(4,24) = 2.844 | 0.046 | Holm-Sidak | OSC vs. SSC<br>SSC vs. NS<br>OSC vs. NS | 0.420<br>0.085<br>0.028 |
|  |  | Central amygdala immunohistochemistry | Two-way RM ANOVA | F(4,24) = 2.844 | 0.046 | Holm-Sidak | OSC vs. SSC<br>SSC vs. NS<br>OSC vs. NS | 0.402<br>0.010<br>0.059 |
|  |  | PVT immunohistochemistry | One-way ANOVA | F(2,12) = 16 | < 0.001 | Tukey | OSC vs. SSC<br>SSC vs. NS<br>OSC vs. NS | 0.681<br><0.001<br>0.002 |
|  |  | Hippocampus subfields immunohistochemistry | Two-way RM ANOVA | F(3,36) = 55.99 | < 0.001 | Holm-Sidak | CA1 vs. DG<br>CA2 vs. DG<br>CA3 vs. DG | <0.001<br><0.001<br><0.001 |
|  |  | CA1 immunohistochemistry | Two-way RM ANOVA | F(6,36) = 2.487 | 0.041 | Holm-Sidak | OSC vs. SSC<br>SSC vs. NS<br>OSC vs. NS | 0.860<br>0.416<br>0.416 |
|  |  | CA2 immunohistochemistry | Two-way RM ANOVA | F(6,36) = 2.487 | 0.041 | Holm-Sidak | OSC vs. SSC<br>SSC vs. NS<br>OSC vs. NS | 0.811<br>0.153<br>0.187 |
|  |  | CA3 immunohistochemistry | Two-way RM ANOVA | F(6,36) = 2.487 | 0.041 | Holm-Sidak | OSC vs. SSC<br>SSC vs. NS<br>OSC vs. NS | 0.968<br>0.465<br>0.465 |
|  |  | DG immunohistochemistry | Two-way RM ANOVA | F(6,36) = 2.487 | 0.041 | Holm-Sidak | OSC vs. SSC<br>SSC vs. NS<br>OSC vs. NS | 0.049<br>0.059<br><0.001 |
| 7 | Day 8 | Fear conditioning freezing and lever presses | One-way ANOVA | $F(14,231) = 0.404$ | 0.973 | n/a | n/a | n/a |
| | | Freezing levels | Kruskal-Wallis | $H(2) = 0.188$ | 0.910 | n/a | n/a | n/a |
| | | Average speed | One-way ANOVA | $F(2,33) = 0.042$ | 0.958 | n/a | n/a | n/a |
| | | Lever press rate | One-way ANOVA | $F(2,33) = 0.480$ | 0.622 | n/a | n/a | n/a |
| | Day 9 | CS lever press rate beginning to end | Paired student's t-test | SE: $t(10) = 1.000$<br>OSE: $t(12) = 1.237$<br>RE: $t(11) = 2.837$ | 0.341<br>0.239<br>0.016 | n/a | n/a | n/a |
| | | Pre-CS lever press rate beginning to end | Paired student's t-test | SE: $t(10) = 1.111$<br>OSE: $t(12) = 3.773$<br>RE: $t(11) = 0.994$ | 0.293<br>0.003<br>0.341 | n/a | n/a | n/a |
| | Day 10 | CS freezing | Two-way RM ANOVA | $F(2,33) = 0.207$ | 0.814 | n/a | n/a | n/a |
| | | CS lever press rate | Two-way RM ANOVA | $F(2,33) = 0.069$ | 0.933 | n/a | n/a | n/a |
| | Day 35 | CS freezing | Two-way RM ANOVA | $F(2,33) = 0.535$ | 0.591 | n/a | n/a | n/a |
| | | CS lever press rate | Two-way RM ANOVA | $F(2,33) = 1.085$ | 0.350 | n/a | n/a | n/a |
| | | pre-CS lever press rate (tone 1) | Two-way RM ANOVA | $F(2,33) = 2.897$ | 0.069 | Holm-Sidak planned comparison | OSE vs. SE<br>OSE vs. RE<br>SE vs. RE | 0.005<br>0.019<br>0.516 |
| | | Average pre-CS lever press rate | Kruskal-Wallis | $H(2) = 6.414$ | 0.041 | Dunn | OSE vs. SE<br>OSE vs. RE<br>SE vs. RE | 0.035<br>0.531<br>0.709 |

On Day 2, rats were placed in context B and given two CS presentations to test the retrieval of tone-associated fear memory in a novel context. Compared to the RC group, both SC and OSC groups exhibited less freezing during the second CS presentation (**Fig. 2C**). Yet, both RC and OSC groups showed reduced average speed during the retrieval test compared to the SC group, suggesting generalized fear responses in a novel context (**Table 1**).

On Day 3, in context A, rats underwent a retrieval and extinction training session with twelve CS presentations. Only the RC group showed a significantly higher pre-tone freezing level, suggesting a robust contextual fear memory (**Table 1**). However, the three groups exhibited the same levels of freezing during the first two CS presentations, suggesting tone-evoked fear retrieval was comparable among the groups (**Fig. 2C**). Nevertheless, a within session extinction analysis comparing the first four CSs to the last four CSs revealed impaired extinction learning in the OSC group as compared to the SC group, with the OSC group displaying smaller reduction of freezing levels from the first to the last CS presentation (**Fig. 2D**). Despite this extinction impairment, no significant differences were found between the OSC and the SC groups during extinction retrieval tests performed on Days 4 and 5 or the spontaneous recovery tests performed on Days 29 and 30 (**Fig. 2C**). The RC group showed higher averaged CS freezing compared to SC group during both extinction retrieval tests (**Table 1**).

In summary, the OSC protocol demonstrated modest enhancement of fear acquisition, indicated by reduced locomotion and stronger, more resistant CS-evoked fear memory during within-session extinction. However, this enhancement didn’t persist during extinction retrieval and spontaneous recovery tests. These results suggest OSC produces a more robust initial fear memory but without a lasting effect on extinction memory.

### The optimal short conditioning protocol induces stronger fear memory than a spaced short conditioning protocol

Spaced learning protocols result in stronger memories than massed learning protocols in both humans (51, 52) and rodents (53-55, but also see 56), raising the possibility that the augmented fear memory observed in the OSC group could simply be attributed to a trial-spacing effect. Our simulations predicted a broad range of effective simulated 4-trial protocols (**Fig. 3**). One protocol with relatively long, fixed ITIs of 11 min 10 s produced a nearly identical peak level of *inducer* as the OSC protocol, and a higher peak level of *inducer* than other protocols with fixed ITIs from 1 min to 21 min (**Fig. 3A**, **Fig. 4A 1**). We termed this protocol the Spaced Short Conditioning (SSC) protocol. Further simulations with reduced stimulus intensity showed that the OSC protocol still produced the greatest peak level of *inducer* and was relatively more effective compared to SSC, with these weaker stimuli (**Fig. 3B**, **Fig. 4A 2**). Therefore, we predicted that the irregular intervals of OSC may produce more effective conditioning than does regularly spaced SSC.

**Figure 3.**
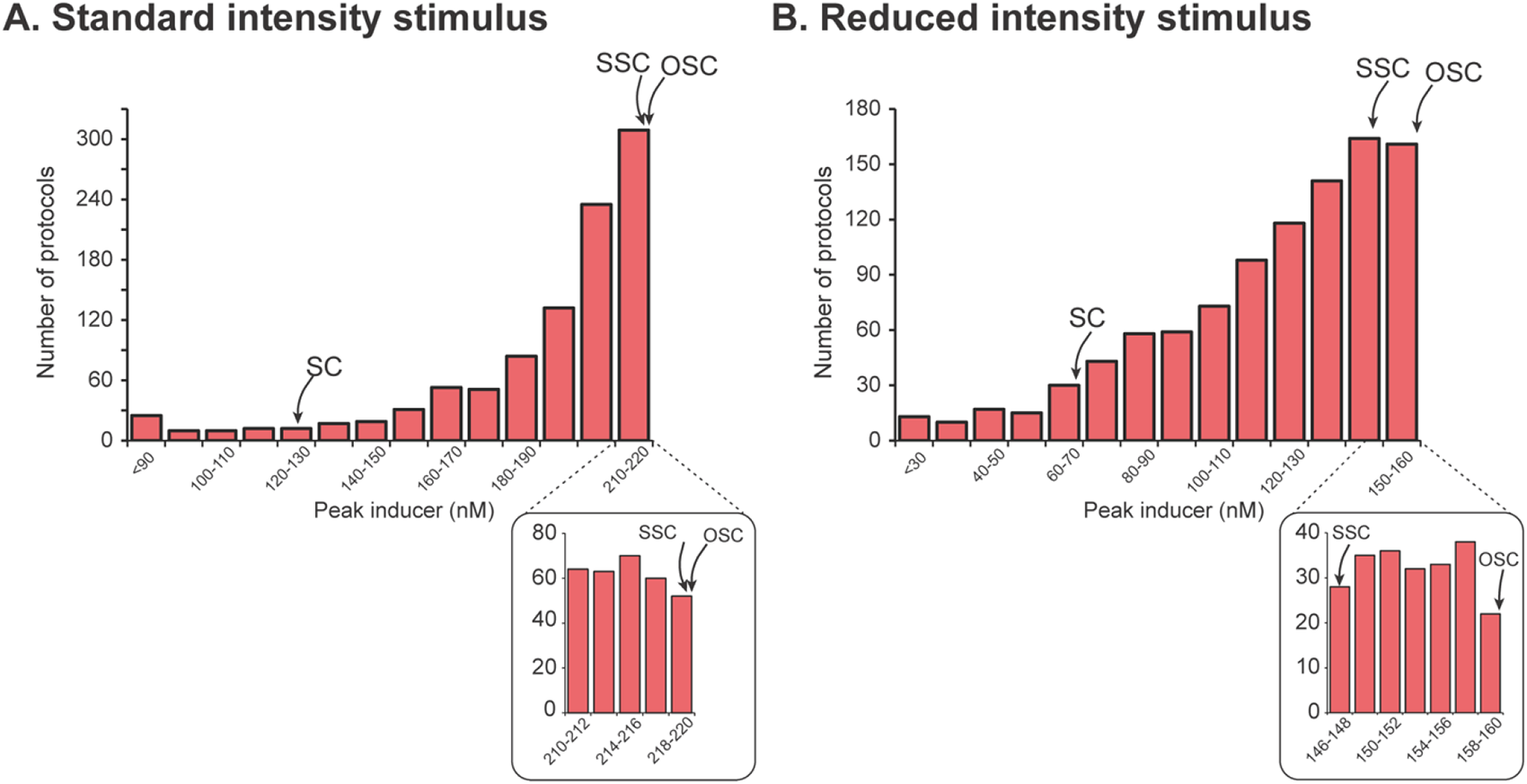
Histogram of peak levels of *inducer* from 1,000 protocols. **A**, Standard intensity stimulus (*Stim* = 300 μM). The range of peak levels of *inducer* (0 – 220 nM) was subdivided into 14 bins, and the number of simulations that produced a peak concentration of *inducer* in each subdivision was plotted. The arrows indicate which bins contained the peak concentrations produced by the short conditioning (SC), spaced short conditioning (SSC), and optimal short conditioning (OSC) protocols. **B,** Reduced intensity stimulus (*Stim* = 200 μM). The range of peak levels of *inducer* (0 – 160 nM) was subdivided into 14 bins, and the number of simulations that produced a peak concentration of *inducer* in each subdivision was plotted. Insets below the main plots illustrate in detail the difference in peak *inducer* for SSC *vs*. OSC, which is negligible in **A**.

**Figure 4.**
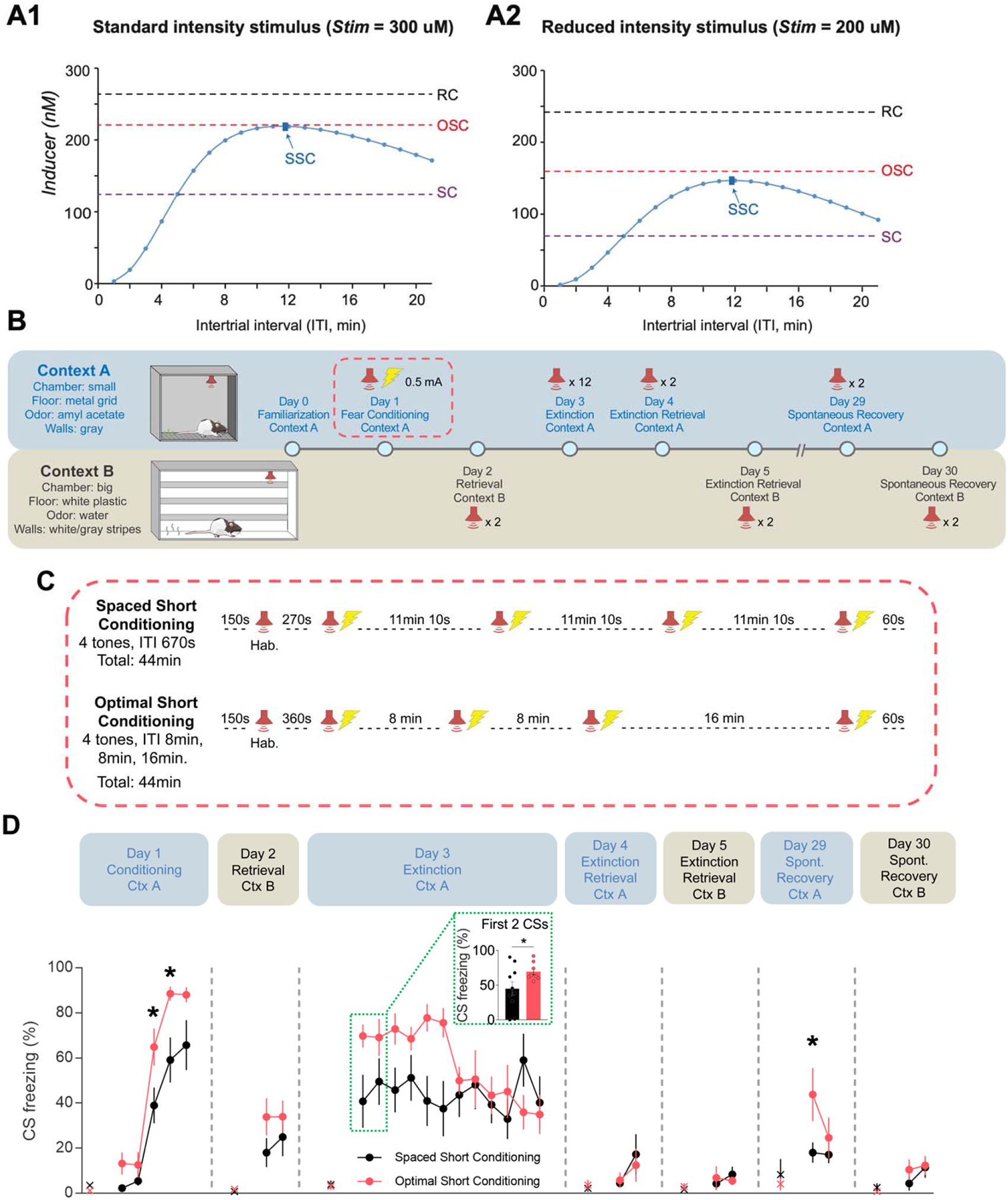
Optimal short conditioning protocol induced stronger fear memory than spaced short conditioning protocol in rats. **A**, *Inducer* peak levels from the RC, SC, and OSC protocols, compared to peak levels from short conditioning protocols of 4 trials with regular ITIs varying from 1 to 21 min, using standard intensity stimulus (**A1**) or reduced intensity stimulus (**A2**). The peak *inducer* values of RC, SC, and OSC are labeled by dashed lines (black: RC; red: OSC; purple: SC). The blue curve gives peak *inducer* values for the protocols with regular ITIs, and the curve peaks at the dark blue dot and arrow, representing the SSC protocol with equal ITIs of 11 min 10s. **B,** Schematics of the fear conditioning procedures. Top: tests that were conducted in Context A. Bottom: tests that were conducted in Context B. **C,** Schematics of the fear conditioning protocols for SSC (n = 10, ITI = 11 min 10 s) and OSC (n = 8) groups. Following a habituation (Hab.) tone, rats received 4 trials of a CS (3 kHz tone, 30 s) that co-terminated with a US (footshock, 0.5 mA, 0.5 s). **D,** Freezing levels during CS presentations of each group across the experiment; ‘X’ denotes the pre-tone freezing levels. Two-way repeated-measure ANOVA for each day followed by Holm-Sidak’s pairwise post hoc test, * p<0.05. Inset: the average freezing level during the first two CS presentations; Welch’s t-test, * p<0.05.

To test this prediction, we empirically compared the SSC and OSC protocols using a reduced shock intensity (0.5 mA instead of the 0.7 mA used in Fig. 2) (**Fig. 4B-C**). Rats exposed to OSC showed higher CS freezing during the second and third CS-US pairing of the fear acquisition session (**Fig. 4D, Day 1)**. In addition, fear memory retrieval was increased when rats in the OSC group were returned to context A for an extinction training session on Day 3, as indicated by higher freezing during the first two CS presentations, compared to the SSC group (**Fig. 4D, Inset**). Both OSC and SSC groups exhibited the same levels of freezing by the end of the extinction training session (**Fig. 4D, Day 3**), as well as during the extinction retrieval tests performed in context A (**Fig. 4D, Day 4**) or context B (**Fig. 4D, Day 5**). However, the OSC group showed greater spontaneous recovery of fear memory in context A approximately 3 weeks later, as indicated by higher freezing during the first CS presentation compared to SSC-trained rats (**Fig. 4D, Day 29**). In summary, these data suggest the enhancement in fear memory acquisition observed with our OSC protocol cannot simply be attributed to a trial-spacing effect or differences in the delay to remove the animals from the chamber, but it is rather associated with the maximized overlap between PKA and ERK signaling.

### The optimal short conditioning protocol induces greater levels of phosphorylated CREB in DG than a spaced short conditioning protocol

The computational model predicts that the increased overlap of PKA and ERK activation in OSC protocol leads to greater LTP induction (*inducer*) in rat hippocampus. This increased overlap plausibly augments phosphorylation of CREB (29), a transcription factor required for synaptic plasticity and LTP (22, 57, See review by 58). We therefore used immunohistochemistry to quantify levels of pCREB following fear conditioning in No-Shock control (NS), and in SSC and OSC groups (**Fig. 5A**) in brain regions previously implicated in the acquisition of fear memory such as the amygdala, the paraventricular nucleus of the thalamus, and the dorsal hippocampus (59, 60, see review by 61). During the fear acquisition session, rats exposed to SSC and OSC protocols exhibited higher freezing levels, indicative of successful fear acquisition compared to the NS group, with the OSC group showing even higher freezing levels than the SSC group (**Fig. 5B**), consistent with Fig. 4.

**Figure 5.**
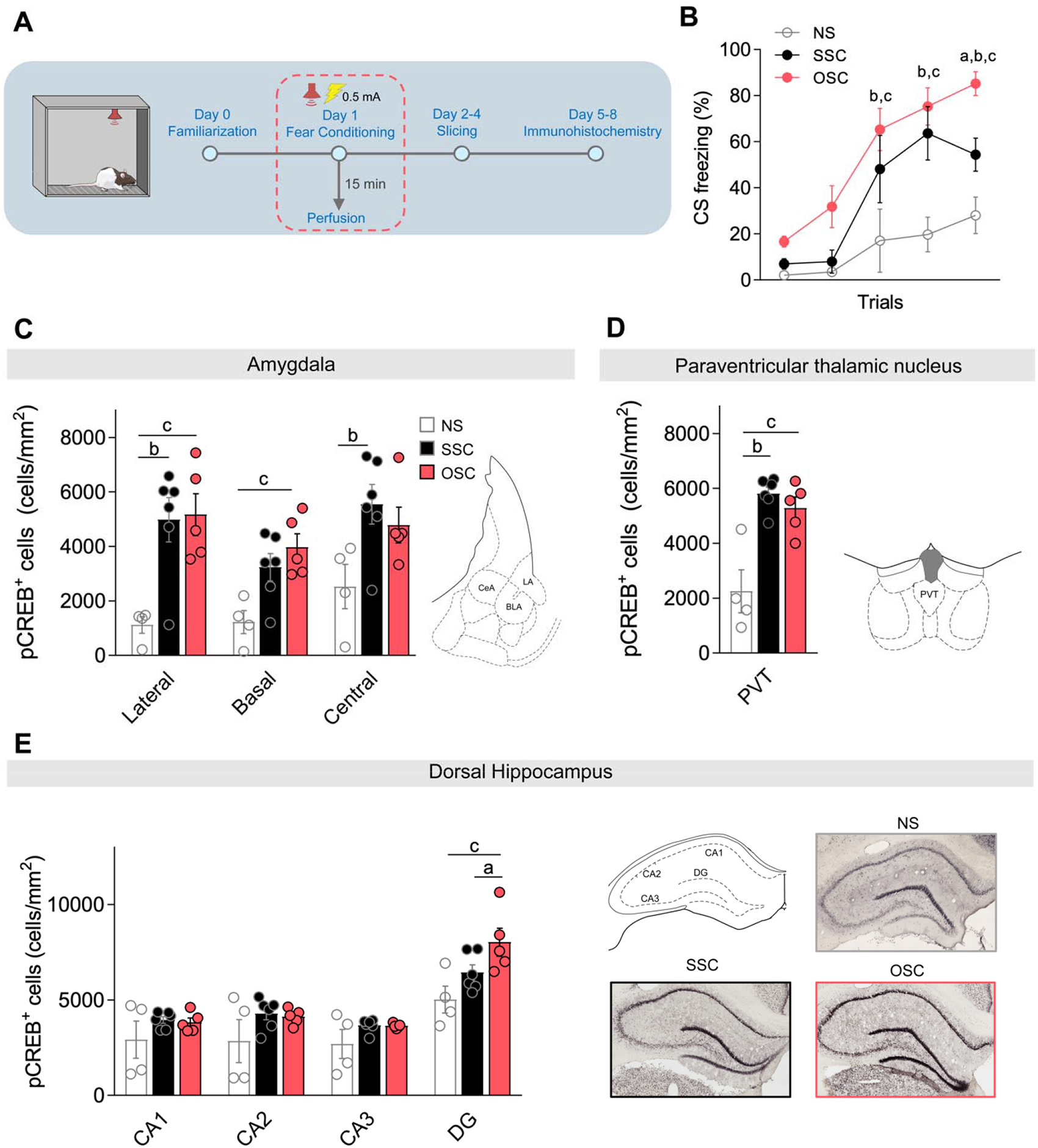
Immunohistochemistry quantification of pCREB in rats exposed to optimal and spaced short conditioning protocols. **A,** Schematic of the immunohistochemistry procedure. **B,** Freezing levels during CS presentation for No-Shock control (NS, n = 4), Spaced Short Conditioning (SSC, n = 6), and Optimal Short Conditioning (OSC, n = 5) groups during fear conditioning; Two-way RM ANOVA followed by Holm-Sidak’s post-hoc test. Letters *a, b*, and *c* indicate pairwise post hoc tests with p < 0.05: *a*, OSC vs. SSC; *b*, SSC vs. NS; *c*, OSC vs. NS. **C,** Average pCREB density in the lateral (LA), basolateral (BLA), and central (CeA) subregions of the amygdala for NS, SSC, and OSC groups; Two-way RM ANOVA followed by Holm-Sidak’s post-hoc test. Letters *b* and *c* indicate pairwise post hoc tests with p < 0.05: *b*, SSC vs. NS; *c*, OSC vs. NS. **D,** Average pCREB density in the paraventricular nucleus of the thalamus (PVT) for NS, SSC, and OSC groups; One-way ANOVA followed by Tukey’s post-hoc test. Letters *b* and *c* indicate pairwise post hoc tests with p < 0.05: *b*, SSC vs. NS; *c*, OSC vs. NS. **E, *Left*,** Average pCREB density across hippocampal subfields for NS, SSC, and OSC groups; Two-way RM ANOVA followed by Holm-Sidak’s post-hoc test. Letters *a* and *c* indicate pairwise post hoc tests with p < 0.05: *a*, OSC vs. SSC; *c*, OSC vs. NS**; *Right***, Representative images of pCREB immunostaining in distinct subfields of the dorsal hippocampus (*top left*) in the NS group (*top right*), SSC group (*bottom left*), and OSC group (*bottom right*).

Immunohistochemistry revealed high levels of pCREB expression in the SSC and OSC groups, as compared to the NS group, across different brain regions including the lateral, basal, and central nuclei of the amygdala (**Fig. 5C**), and the paraventricular nucleus of the thalamus (PVT, **Fig. 5D**), suggesting that increased pCREB levels in SSC and OSC groups are mediated by the CS-US pairing. However, we did not observe differences in pCREB levels between the OSC and SSC groups in the amygdala or PVT (**Table 1**), and pCREB levels were similar among the three groups in CA1 hippocampal subfields (**Fig. 5E**). Interestingly, rats exposed to the OSC protocol exhibited increased pCREB expression in the DG, as compared to the SSC and NS groups (**Fig. 5E**). Given that increased pCREB in the DG has been associated with enhanced LTP in rodents (62), these data support the hypothesis that the augmentation in fear memory acquisition observed with the OSC protocol is mediated by higher levels of LTP induction in the rat hippocampus.

### The optimal protocol also enhanced fear extinction

Extinction is a new learning that temporarily inhibits the initial associative memory (63, 64). Extinction also relies on PKA and ERK signaling to induce plasticity-related gene transcription (65–67). We predicted that the computationally designed protocol would also enhance the acquisition of extinction, thereby suppressing the original fear memory (**Fig. 6A-B1**). We compared an Optimal Short Extinction (OSE) protocol, comprising four trials with irregular ITIs of 8, 8, and 16 minutes (**Fig. 6B3**), against a Regular Extinction (RE, **Fig. 6B4**) protocol using 12 trials and ITIs of 150 s, and a Short Extinction (SE, **Fig. 6B2**) protocol using 4 trials and the same ITIs of 150 s, which resemble previous protocols used for regular and short fear extinction in rats (68–70). In simulations, the OSE protocol triggered higher peak *inducer* than the SE protocol, and was comparable to the RE protocol (**Fig. 6B1-3**). We thus predicted OSE would result in stronger extinction of fear memory than does the standard SE.

**Figure 6.**
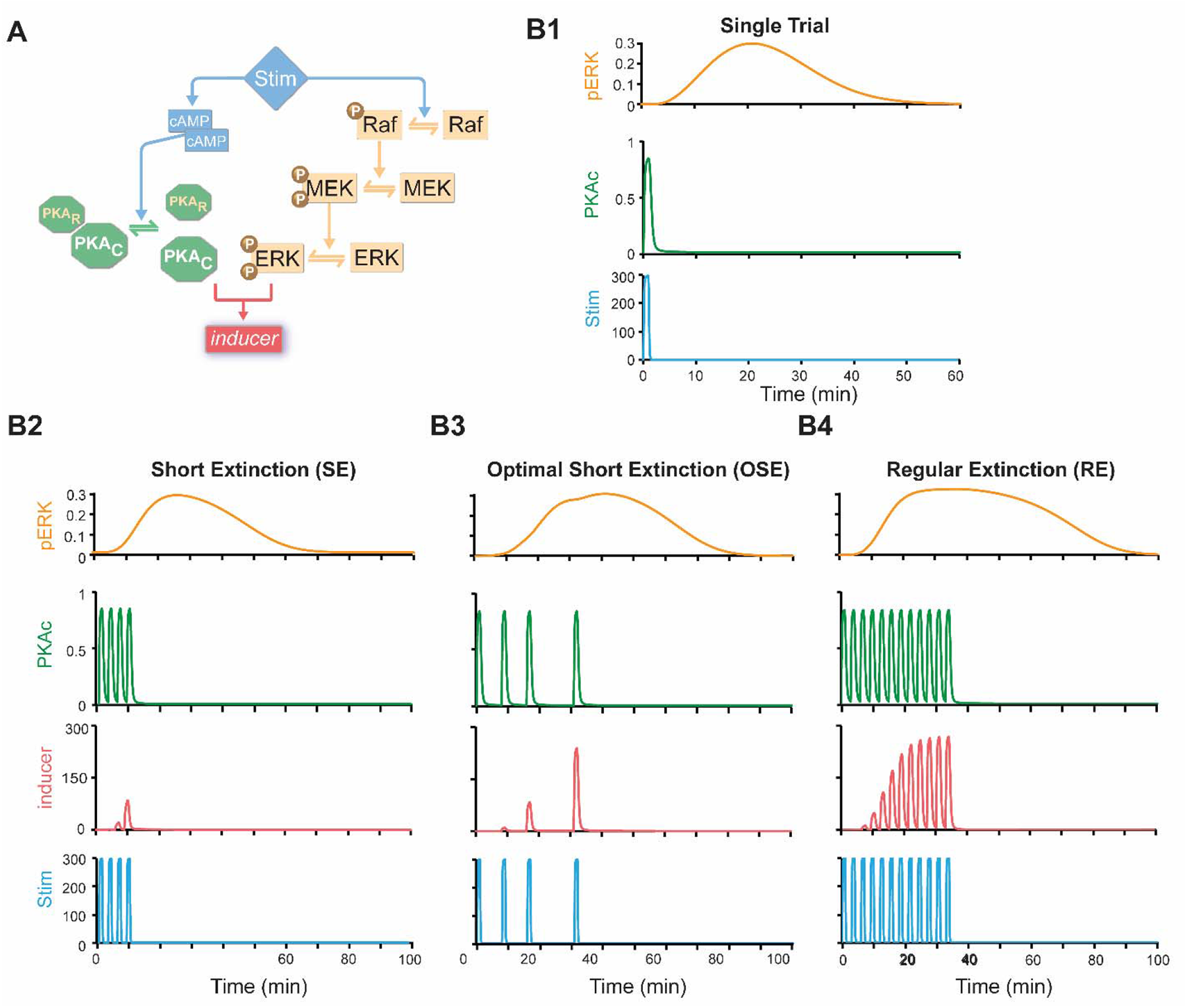
Prediction of enhanced protocol for fear extinction. **A**, Schematic of the model. **B1**, Simulated time courses of activated ERK (pERK, μM) and activated PKA (PKAc, μM) in response to one trial of *Stim* (μM). **B2**, Simulated time courses of pERK, PKAc, and *inducer* in response to 4-trial protocol with regular intervals of 2.5 min (short extinction, SE). **B3**, Simulated time courses of pERK, PKAc, and *inducer* in response to 4-trial protocol with computationally designed intervals (optimal short extinction, OSE). **B4**, Simulated time courses of pERK, PKAc, and *inducer* (nM) in response to 12-trial protocol with regular intervals of 2.5 min (regular extinction, RE).

To test this prediction, we designed an experiment comparing extinction protocol efficacy using conditioned suppression of reward-seeking behavior (**Fig. 7A**), which is more sensitive than freezing in fear extinction paradigms (71, 72). Lever presses also help to maintain constant activity for reliable freezing measurement along the session (73). Since lever presses for reward are trained in a specific context, and extinction memory is context-dependent (74, 75), we used only one context in this experiment. Rats were initially trained to press a lever to receive sucrose pellets in a variable interval schedule of 60 s until they reached the same levels of lever pressing and locomotor activity (see Supp. Methods). On Day 8, rats underwent a fear conditioning session which included five nonreinforced habituation tones followed by seven CS-US pairings. On Day 9, rats were pre-assigned to three experimental groups for extinction (**Fig. 7B**) so that freezing levels, average speeds, and lever-press rates were similar during the fear conditioning session (**Table 1**). At the end of the extinction session, freezing levels reduced from ∼50% to ∼25% in the RE group, whereas the SE and OSE groups maintained the same levels of freezing throughout the four CSs (**Fig. 7C**, Day 9). Similarly, the RE group showed a significant increase in lever presses during the CS presentations at the end of the session, whereas CS lever presses remained unaltered in the SE and OSE groups (**Fig. 7D *left***). However, the OSE group showed a significant increase in lever presses during the 30-s periods preceding the CS presentations (pre-CS, **Fig. 7D *right***), suggesting enhanced within-session extinction of contextual fear memory (76).

**Figure 7.**
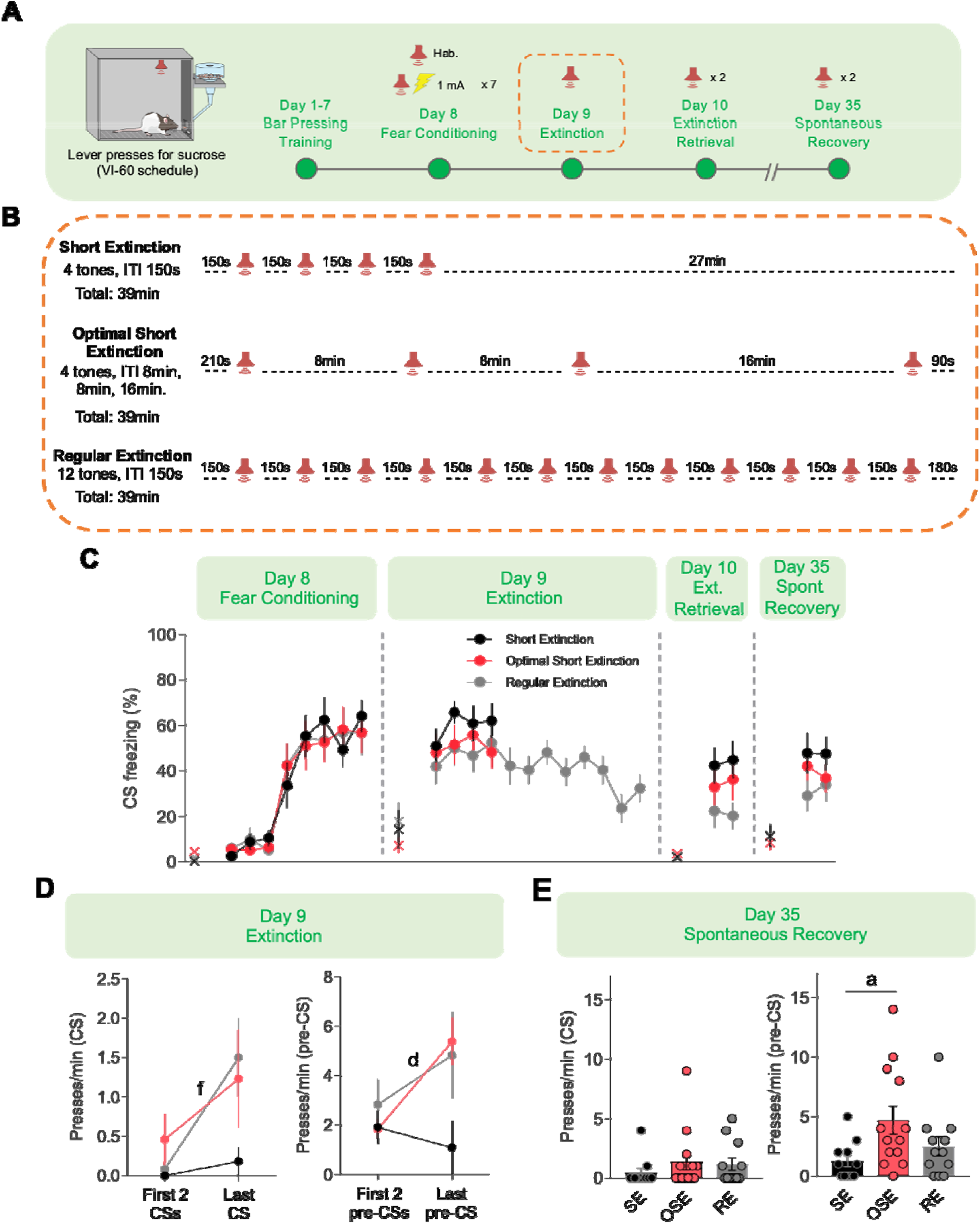
The computationally designed protocol partially enhanced fear extinction in rats. **A.** Schematic of the enhanced fear extinction experimental procedures. Rats were trained to constantly press a lever to retrieve sucrose pellets in a variable interval 60 (VI60) schedule, where the average interval between each sucrose delivery is 60 s. **B,** Schematics of the fear extinction protocols for SE (n = 11), OSE (n = 13), and FE (n = 12) groups, during which rats received multiple trials of a CS (3 kHz tone, 30 s). **C,** Freezing levels during CS presentations of each group across the experiment. No significant difference between groups was found by Two-way RM ANOVA. ‘X’ denotes the pre-tone freezing levels. **D,** Lever press rates during the first two and the last CS presentation (*left*) or the 30 s before the first two and the last CS presentation (pre-CS period) of each group. OSE group shows significant increase of pre-CS lever presses comparing the last CS presentation to the average lever press rate of the first two CS presentations. Paired Student’s t-test. Letters *f* and *d* indicate tests with p < 0.05: *f*, RE, Last CS vs. First 2 CS; *d*, OSE, Last pre-CS vs. First 2 pre-CS. **E,** OSE group shows higher lever presses during pre-CS period compared to SE group in spontaneous recovery test. No difference was found for the CS lever presses among the three groups (*H(2)* = 2.102, *P* = 0.349). Kruskal-Wallis’s test followed by Dunn’s post hoc test. Letter *a* indicates pairwise comparison with *p<0.05: OSE vs. SE.

On Days 10 and 35, rats were returned to the same chamber to test the strength of fear extinction memory during extinction retrieval and spontaneous recovery tests, respectively. No significant difference in CS freezing or CS lever presses were observed among the three groups (**Fig. 7C, Day 10&35**). However, the OSE group showed increased pre-CS lever pressing during the spontaneous recovery test (**Fig. 7E *right***), when compared to the SE and RE groups. These results suggest that, although the tone-associated memory was similar among the groups, the OSE protocol enhanced contextual fear memory extinction, evidenced by enhanced within-session extinction of conditioned suppression and reduced conditioned suppression three weeks post-extinction training.

## DISCUSSION

A computationally designed protocol that maximizes the interaction between PKA and ERK pathways enhances LTF and nonassociative learning in *Aplysia* (26). Here, we extended this strategy to associative learning in mammals by adapting the simplified mathematical model used in *Aplysia* to simulate the dynamics of PKA and ERK in rat hippocampus based on empirical data in the literature (33–35). Through a series of computational simulations and experimental validations, we discovered that an optimized training protocol, predicted to enhance the overlap of PKA and ERK activation, can facilitate acquisition and extinction of conditioned fear. Immunohistochemistry demonstrated that the augmented fear memory induced by the optimal protocol was associated with increased expression levels of pCREB in the DG subfield of the dorsal hippocampus. Our results demonstrate the power of a simplified model of intracellular signaling cascades in describing associative learning across species, attesting the essential role of the interaction between PKA and ERK pathways in both nonassociative and associative learning.

Training protocols using spaced ITIs result in stronger memory acquisition than those using standard massed ITIs, a well-established phenomenon described as the “spacing effect” (see review by 11). The spacing effect has been linked to the varying efficacy of massed and spaced protocols in triggering LTP/LTF via biochemical signaling, and a previous computational model suggests the optimal ITI aligns with peak ERK activation (77–79). However, that model assumed fixed ITIs, which may not be ideal for synaptic plasticity and long-term memory. Our optimized training protocol, with irregular ITIs, induced stronger and persistent fear memory in rats, as compared to a spaced protocol that had been predicted to be the most effective among protocols with equal ITIs. These results suggest that the spacing effect in mammals can be at least partially explained by enhanced overlap between the PKA and ERK pathways, which are critical for CREB activation (29). Our study also indicates that learning protocols with irregular inter-trial intervals (ITIs) may be more effective than those with equal ITIs. Although spaced protocols are known for facilitating fear memory acquisition, there is a lot of controversy when it comes to fear extinction. Whereas some studies have demonstrated that spaced intervals between the CSs facilitate extinction memory and attenuate renewal and spontaneous recovery of fear (56, 80–82), others have shown the opposite, impaired extinction memory and increased recovery of fear (56, 81) . We therefore selected a control group with massed trials to better control for our optimized protocol.

The enhanced performance of the optimal (OSC) protocol appeared to be constrained by the intensity of the stimuli. The model predicted higher performance when weak stimuli were used but comparable performance with strong stimuli (**Fig. 4A**). Drawing a line to distinguish weak and strong footshock intensities is not straightforward because the relationship between fear memory and footshock intensity is neither monotonic nor linear (83). Nevertheless, we observed clear differences in the efficacy of the optimal fear conditioning protocol when using footshocks of different intensities. When we compared the OSC protocol to a massed short conditioning protocol using a standard footshock intensity (0.7 mA, **Fig. 2**) commonly used in previous studies (28), we found only small differences in the conditioned responses (CS freezing) between the two protocols. However, when we used a lower footshock intensity (0.5 mA, **Fig. 4**) to compare the optimal short conditioning protocol to a spaced short conditioning protocol, we found a significant increase in CS freezing in the OSC group during fear acquisition, retrieval, and spontaneous recovery tests. Considering the robustness of the spacing effect, it is possible that differences between the OSC and the massed SC groups at the higher footshock intensity were in part masked by a “ceiling effect”. Alternatively, previous studies in rats have demonstrated that the hippocampus is required for tone-evoked fear memory when a weak (0.4 mA) but not a strong (0.9 mA) footshock is used (84), which could also explain the differences we observed at the two intensities. Either way, our data suggest that our approach could be more beneficial for learning protocols relying on relatively weak stimuli. Additional studies need to be conducted to understand the mechanism by which the intensity of stimuli governs the enhanced performance of protocols with irregular ITIs compared to protocols with fixed ITIs.

The association between the CS and US is primarily mediated by the lateral amygdala where LTP induces enhanced CS responses (see review by 39, 85). Nevertheless, another important and distinguishable component of fear memory formation is the context in which the association has occurred. Interestingly, we only observed enhanced conditioned responses in the context in which fear conditioning occurred, suggesting the memory facilitation effect is context-dependent. Similarly, in the fear extinction experiment, enhancement by the optimal protocol was only observed during the pre-CS lever pressing, a more sensitive index of contextual fear memory during extinction and spontaneous recovery (71, 72, 76, 86, 87). This is consistent with the hippocampus’ role in encoding context during fear conditioning and extinction, and in the time-dependent reappearance of fear post-extinction (*i.e.*, spontaneous recovery), on which our model is based (see reviews by 46, 88). Accordingly, immunohistochemical results showed greater pCREB expression in the OSC group specifically in the DG of the dorsal hippocampus, a region implicated in the acquisition and extinction of contextual fear memory (89, 90). Increasing CREB expression in DG neurons has been demonstrated to improve contextual fear acquisition (91), whereas inactivating DG neurons that overexpress CREB disrupts contextual fear retrieval (92).

Another hippocampal subregion implicated in contextual fear is the CA1 (93). Although the OSC, SSC and NC groups displayed similar levels of pCREB expression in CA1, the single post-training time point (15 min) we used does not suffice to rule out the possibility of distinct pCREB dynamics in CA1 compared to DG. CA1 pCREB peaks at 30 min and disappears at 90 min following fear conditioning (94, 95). Further experiments comparing distinct post-training intervals may better reveal the dynamics of pCREB expression in different hippocampal subregions, as well as in other areas involved in fear acquisition (e.g., amygdala, PVT), which in the present study exhibited similar levels of pCREB expression in the OSC and SSC groups. The observation that the dynamics of ERK pathways differ across brain regions involved in fear memory, with peaks occurring at 20 min after a single stimulation in the hippocampus versus 60 min in the lateral amygdala (35, 96, 97), suggests that the model may be able to predict optimal protocols targeting specific components of fear memory based on the dynamics of intracellular signaling cascades in corresponding brain regions. Future studies will test this possibility by modeling the molecular cascades in the lateral amygdala and the medial prefrontal cortex to preferentially target the acquisition and extinction of CS-associated memories, respectively.

Our model is an obvious simplification of the mechanisms underlying fear conditioning and extinction. It does not include other molecular cascades critical for LTP and memory formation, such as calcium/calmodulin-dependent kinase II (CaMKII) or protein kinase C (PKC), due to the lack of empirical data for simulating their dynamics of activation (12, 98, 99). The model was constructed using empirical PKA and ERK dynamics from the literature, which are based on *ex vivo* analyses and have limited temporal resolution. Furthermore, the model assumes the molecular mechanisms for fear memory acquisition are similar to those for extinction. Despite these limitations, the simplified model was sufficient to predict enhanced activation of the LTP-related transcription factor CREB in the DG and to facilitate associative learning in three different experiments. A more complex model that incorporates a wider range of intracellular and extracellular processes based on *in vivo* data would likely have enhanced predictive ability. Further experiments should also investigate the memory enhancing effects of computationally designed training protocols in different types of associative memories, including discrimination learning and backwards conditioning. In addition, it will be important to test whether the effects observed with optimal protocols vary across subjects of different sexes and ages, as well as protocol efficacy in animal models of cognitive impairment. Together, our results suggest the possibility of using similar model-driven, non-invasive behavioral approaches in studies aimed at enhancing learning or restoring memory deficits in humans, or aimed at improving extinction-based exposure therapies for anxiety disorders.

## MATERIALS AND METHODS

### Animals

The study involved 120 male Long-Evans rats, aged 3-5 months, kept in a 12-hour light/dark cycle. All procedures followed the National Institutes of Health guidelines for animal care and were approved by the Center for Laboratory Animal Medicine and Care of The University of Texas Health Science Center at Houston.

### Model Development

The mathematical model, adapted from (26), describes the activation of the PKA and ERK pathways. Parameters were adjusted based on empirical data (33–35, 78) for hippocampus. Simulations used fourth-order Runge-Kutta integration for differential equations, and were conducted in XPPAUT (100) on Dell Precision T1700 microcomputers.

### Behavioral Tasks

Rats were exposed to fear conditioning in two distinct chambers. Three groups – RC (eight tone-shock pairings, 270s intervals), SC (four pairings, 270s intervals), and OSC (four pairings, 8, 8, and 16 min intervals) – were used. Optimal vs. Spaced Short Conditioning compared OSC with SSC (four pairings, 11min 10s intervals) with footshock intensity at 0.5 mA. Extinction involved lever-press training, followed by fear conditioning. These rats were divided into three groups: RE (twelve tone presentations, 150s intervals), SE (four tone presentations, 150s intervals), and OSE (four tone presentations, 8, 8, and 16 min intervals).

### Immunohistochemistry

Rat brains were prepared post-conditioning for immunohistochemistry to analyze pCREB-positive cells. Brains were sectioned and treated with rabbit anti-pCREB serum (1:1000; EMD Millipore 06-519) and biotinylated goat anti-rabbit IgG antibody (1:200, Vector Labs, BA-1000-1.5), and revealed by ABC kit (1:50, Vector Labs, PK-6100) and DAB-Ni solution (Vector Labs, SK-4100). Images were captured using a Nikon microscope and analyzed with QuPath software (101) for automatic cell detection and quantification. pCREB-positive cells in specified brain regions were counted and densities calculated.

### Quantification and statistical analysis

Grubbs’ tests (102) were used to identify outliers in each experiment. Shapiro-Wilk tests were performed to determine normal distribution. Equal variance was confirmed through F tests and Brown-Forsythe tests. Statistical significance was determined using t-tests, ANOVA, or Kruskal-Wallis tests with relevant post-hoc comparisons, as appropriate. Sample size was based on prior literature and experience.

## Data availability

All data that support the findings presented in this study are available from the corresponding author on reasonable request. Source codes will be submitted to GitHub (https://github.com/Owenxz/Zhang-XO-Enhanced-Learning-2022.git).

## Author Contributions

X.O.Z, D.S.E., and C.E.C performed and analyzed the behavioral experiments. X.O.Z and C.E.C performed and analyzed the immunohistochemistry experiments. Y.Z. implemented the computational model and ran all simulations. P.S. helped design and implement the computational model. F.H.D-M and J.H.B supervised and contributed to all aspects of this study. All the authors participated in the design of the experiments. X.O.Z. and F.H.D-M interpreted the data and prepared the manuscript with comments from all the co-authors.

## Acknowledgments

This work was supported by the Russell and Diana Hawkins Family Foundation Discovery Fellowship and the Dr. John J. Kopchick Fellowship to X.O.Z, NIH grant R01-NS102490 to J.H.B, and NIH grant R00-MH105549, NIH grant R01-MH120136, a Brain & Behavior Research Foundation grant (NARSAD Young Investigator), and a Rising STARs Award from UT System to F.H.D-M. We thank Nikita Watson and Sharon Gordon for their technical and administrative assistance. We also thank current members of the Byrne and Do Monte labs for their valuable comments on the manuscript. This manuscript has been posted on BioRxiv as a preprint (doi: https://doi.org/10.1101/621540).

